# Structural basis for the selective inhibition of the PI3KC3-C2 complex by Rubicon in endolysosome maturation and mitophagy

**DOI:** 10.64898/2026.08.07.743589

**Authors:** Minghao Chen, Aniketh Bishnu, Yongjia Duan, Julia F. Riley, Qingyang Ni, Aaron Joiner, Isaac J. Allen, Erika L. F. Holzbaur, Ian G. Ganley, James H. Hurley

**Author notes:** James H. Hurley **Email:**.

## Abstract

Rubicon is a negative regulator of autophagy and the endolysosomal network (ELN) and an antagonist of the class III phosphatidylinositol 3-kinase complex II (PI3KC3-C2). Inhibition of Rubicon is considered a potential means to therapeutically upregulate autophagy and the ELN to treat Parkinson’s disease and other conditions characterized by autophagic and ELN dysfunction. Rubicon is specific for the UVRAG-containing PI3KC3-C2 over the purely autophagic ATG14- containing PI3KC3-C1 complex. Here, we determined the high-resolution cryo-electron microscopy structure of PI3KC3-C2 in complex with the PI3KC3-binding domain (PIKBD) of Rubicon and compared it to cryo-EM structures of unbound PI3KC3-C2 and PI3KC3-C1. Rubicon binds directly to PI3KC3-C2 only via the BARA domain of the BECN1 subunit, which is common to both C1 and C2. The selectivity of Rubicon for the PI3KC3-C2 complex over the PI3KC3-C1 complex is attributed to a conformation of the BECN1^BARA^ domain induced by UVRAG, rather than to direct contact with UVRAG or direct antagonism by the ATG14 subunit of PI3KC3-C1. Targeted disruption of the Rubicon:PI3K3-C2 structural interface by site-directed mutations enhances mitophagic activity in human epithelial cells to levels comparable to those observed in Rubicon knockout (KO) cells. Similarly, disruption of the interaction in Rubicon-overexpressing hippocampal neurons restored lysosomal flux to wild-type levels. These data show that suppressing the function of PI3K3- C2 can fully account for the negative regulatory effects of Rubicon in the autophagy and ELN pathways.

**Significance Statement:** Endolysosome maturation and autophagosome-lysosome fusion require the production of phosphatidylinositol 3-phosphate (PI(3)P) by the class III phosphatidylinositol 3-kinase complex II (PI3KC3-C2). Rubicon is a key negative regulator of endolysosomes and autophagy that suppresses PI3KC3-C2 activity. Here, we reveal in atomistic detail how Rubicon selectively recognizes PI3KC3-C2. Disrupting the Rubicon–PI3KC3-C2 interaction restores mitophagy and enhances lysosomal activity to the same extent as Rubicon gene deletion, establishing that PI3KC3-C2 inhibition fully accounts for the biological regulatory effects of Rubicon in the autophagy and lysosome pathways.

## Introduction

Macroautophagy (hereafter, “autophagy”) is the eukaryotic cell’s principal means for degrading large materials such as organelles and protein aggregates. Deficits in autophagy are widespread in human diseases associated with the build-up of aggregated proteins and damaged organelles (1, 2). Genetic ablation of autophagy in the brain leads to rapid neurodegeneration in animals (3), which, even at an advanced stage, can be reversed by re-activation of autophagy (4). Autophagy inducers are therefore being sought as potential therapeutics (5). Mechanism-based allosteric activation of core autophagy proteins (6) and phenotypic screening (7) have been among the approaches tried. Inhibition or degradation of negative regulators of autophagy is another approach. Inhibition of the negative regulator mTORC1 is widely used to induce bulk autophagy, but at the cost of also inhibiting many other critical pathways (8). Inhibition of USP30, a negative regulator of PINK1- and Parkin-dependent mitophagy, has shown promise in therapeutic rescue of mitophagy and α-synuclein clearance in mice (9). The best-characterized negative regulator of autophagy is Rubicon (RUN domain Beclin-1-interacting and cysteine-rich containing protein (10–12). Knockout of Rubicon in mice has favorable properties, including reduced α-synuclein aggregation, with no negative effects noted other than male sterility (13). Recent work identified Rubicon as a negative regulator of mitophagy (14) in neurons subjected to oxidative stress. These data make Rubicon an attractive target for autophagy-enhancing therapeutics and motivated this study of its mechanism of action.

Rubicon was discovered based on its interaction with the class III phosphatidylinositol 3-kinase complex II (PI3KC3-C2) (10–12). PI3KC3-C2 is a lipid kinase complex that plays essential roles in endolysosome maturation by producing phosphatidylinositol 3-phosphate (PI(3)P) (15–21). PI3KC3-C2 consists of four subunits, one copy each of the catalytic lipid kinase VPS34, the scaffold protein VPS15, and two regulatory subunits, Beclin-1 (BECN1) and UV radiation resistance-associated gene protein (UVRAG) (22–24). Rubicon belongs to the Rubicon Homology (RH) domain-containing protein family, which also contains Pacer (25, 26) and PLEKHM1 (27). Rubicon is a 972-amino acid protein that can be subdivided into three regions. The N-terminal RPIP8, UNC- 14, and NESCA (RUN) domain is important for biological function (28), but its precise biochemical role is unknown. The central intrinsically disordered region (middle region, MR) contains within it a PI3K-binding domain (PIKBD) (23). The C-terminal Rab7-binding RH domain has a complex multiple Cys-Zn cluster fold and is responsible for endolysosomal localization (27, 29, 30).

PI3KC3-C2 is important for the broad functioning of the endolysosomal pathway, including autophagy, while the related complex PI3KC3-C1 is uniquely specialized for autophagy. PI3KC3- C1 differs from C2 only in that the UVRAG subunit is replaced by the autophagy-specific subunit ATG14 (11, 15, 21, 31). UVRAG and ATG14 integrate into their respective PIKC3 complexes by forming a parallel coiled-coil dimer with BECN1 (32, 33). The presence of UVRAG or ATG14 is mutually exclusive because the BECN1 coiled-coil only possesses one interface capable of binding these proteins. Both UVRAG and ATG14 contain BARA-like domains (UVRAG BARA2 domain (residues 327-461); ATG14 C-terminal domain (CTD, residues 206-412)) C-terminal to their coiled coil. The BARA-like domains contact the BARA domain of BECN1. The BECN1 BARA domain is a major determinant of PI3KC3 docking to membranes (34, 35). PI3KC3-C1 is targeted to membranes by RAB1A (24) while C2 is targeted by RAB5A (24, 36, 37). PI3KC3-C1, but not C2, forms a supercomplex with the other major autophagy-initiating complex, the ULK1 complex (38), which together comprise the core of the mammalian autophagy initiation machinery (39, 40). Rubicon binds to PI3KC3-C2 but not C1. We previously reported an intermediate resolution structure of a Rubicon-PI3KC3-C2 complex (23). This structure did not resolve side-chain level detail, however, and was insufficient to rationalize the preference of Rubicon for C2 over C1. Thus, the signature characteristic of Rubicon, its selective inhibition of the C2 but not C1 complex, still needed to be explained at the structural level.

Here, we elucidate the mechanism underlying the selective binding by determining cryo-EM structures of PI3KC3-C2 in complex with the Rubicon^PIKBD^ region, as well as the unbound forms of PI3KC3-C2 and C1, at resolutions of 3.38 Å, 3.83 Å, and 3.77 Å, respectively. Comparative structural analyses reveal that the Rubicon binding site within the BECN1^BARA^ domain undergoes a pronounced conformational rearrangement governed by the presence of either the UVRAG^BARA2^ domain or ATG14^CTD^ domain, thereby indirectly modulating Rubicon binding specificity. We take advantage of structural insights to surgically cripple PI3KC3-C2 binding and assess the impact on mitophagy and lysosomal function.

## Results

### Structural mapping of the Rubicon:PI3KC3-C2 interface

The Rubicon:PI3KC3-C2 complex measures approximately 225 × 135 × 70 Å and adopts an L-shaped overall architecture comprising a catalytic arm, an adaptor arm, and a base region (Fig. 1A-B), like other PI3KC3 structures. All the domains of PI3KC3-C2 were well resolved in the present structure, except for the VPS34 kinase domain. Focused refinement of the adaptor arm region in the Rubicon:PI3KC3-C2 sample yielded a local resolution at 3.38 Å, enabling accurate model building with confident side-chain assignment (SI Appendix Fig. S1). An additional EM density was observed adjacent to the BECN1^BARA^ domain and was assigned to the helix α1 of the Rubicon^PIKBD^ (residues 488-515) (SI Appendix Fig. S2A). This assignment is consistent with our previous finding that helix α1 is the minimum element required for interaction with PI3KC3-C2 (23). The BECN1^BARA^ domain in PI3KC3-C2 forms a highly complementary surface that accommodates the Rubicon^PIKBD^ helix α1 (Fig. 1C-D, SI Appendix Fig. S2B). Specifically, the Rubicon-binding interface comprises a positively charged groove flanked by two hydrophobic pockets that engage the negatively charged E492, S499, and E507 and the hydrophobic side chains of F496 and I503 on helix α1, respectively (Fig. 1D-E, SI Appendix Fig. S2B).

**Figure 1.**
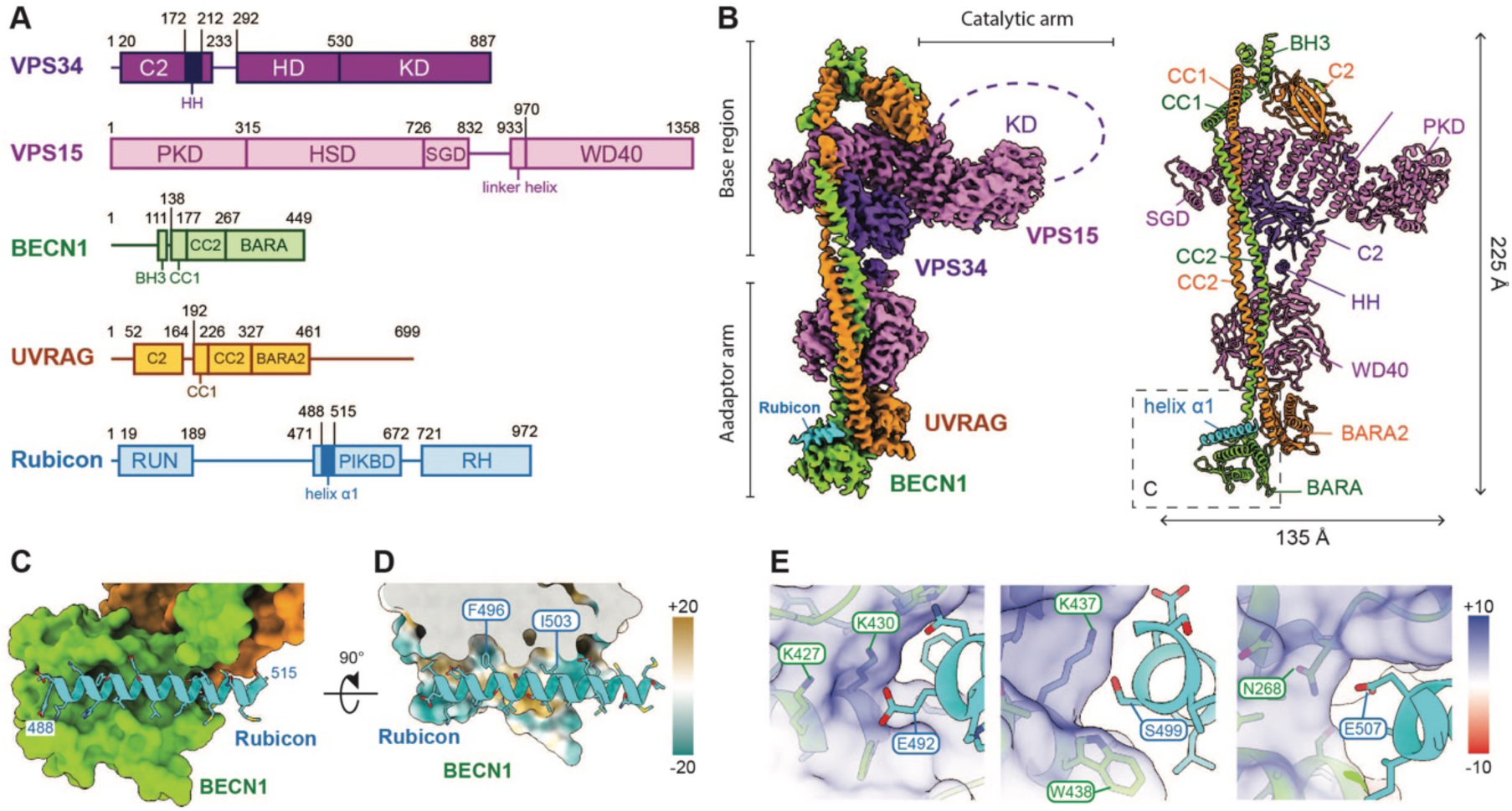
Cryo-EM structure of the Rubicon:PI3KC3-C2 complex. (A) Domain organization of PI3KC3-C2 and Rubicon. (B) Overall cryo-EM map (left) and the corresponding atomic model (right) of the Rubicon complex. Subunits and domains are labeled and colored as in A. (C) Close-up view of the Rubicon-binding site on the BECN1 BARA domain. PI3KC3-C2 is shown as a surface, and the Rubicon α1 helix as a ribbon. (D) Cross-sectional view of the Rubicon-binding site. The view is rotated by 90° relative to C to illustrate the surface complementarity. The PI3KC3-C2 surface is colored according to molecular lipophilicity potential (cyan, −20; yellow, +20), and the two key hydrophobic residues of Rubicon, F496 and I503, are indicated. (E) Close-up view of the electrostatic interactions mediating Rubicon binding. The BECN1 surface is displayed as a transparent surface colored by electrostatic potential (red, −10; blue, +10). Residues involved in the interaction are indicated.

To validate the identified interface, we designed two sets of mutations (Mut1: E492K/S499K/E507K and Mut2: F496D/I503D) targeting either electrostatic or hydrophobic interactions. Bead-binding assays revealed a dramatic loss of binding to the Rubicon fragment (478-512) upon the introduction of either mutation or upon their combination, confirming that these residues are critical for the Rubicon-PI3KC3-C2 interaction (Fig. 2A-B). Since the effects on PI3KC3-C2 binding were the same, we treated these mutations as equivalent perturbations in subsequent experiments. In contrast to PI3KC3-C2, but consistent with previous findings (10–12), no interaction was observed between Rubicon and PI3KC3-C1. A complementary PI3KC3-C2 bead-binding experiment in which the bait and prey roles were switched yielded the same results, validating the significance of the proposed residues (SI Appendix Fig. S3). We further tested how this interaction influences PI3KC3- C2 activity by measuring ATP consumption in the presence of wild-type (WT) or mutant Rubicon fragments, using small unilamellar vesicles (SUVs) as membrane substrates. The WT fragment markedly inhibited PI3KC3-C2 activity, whereas the quintuple mutant (Mut1+2: E492K/S499K/E507K/F496D/I503D), which disrupts the Rubicon-BECN1 interface, largely restored ATP consumption to a level comparable to that observed in the absence of the Rubicon fragment (Fig. 2C). These results demonstrated that binding of the Rubicon helix α1 is necessary for the inhibition of PI3KC3-C2 activity in an *in vitro* biochemical assay.

**Figure 2.**
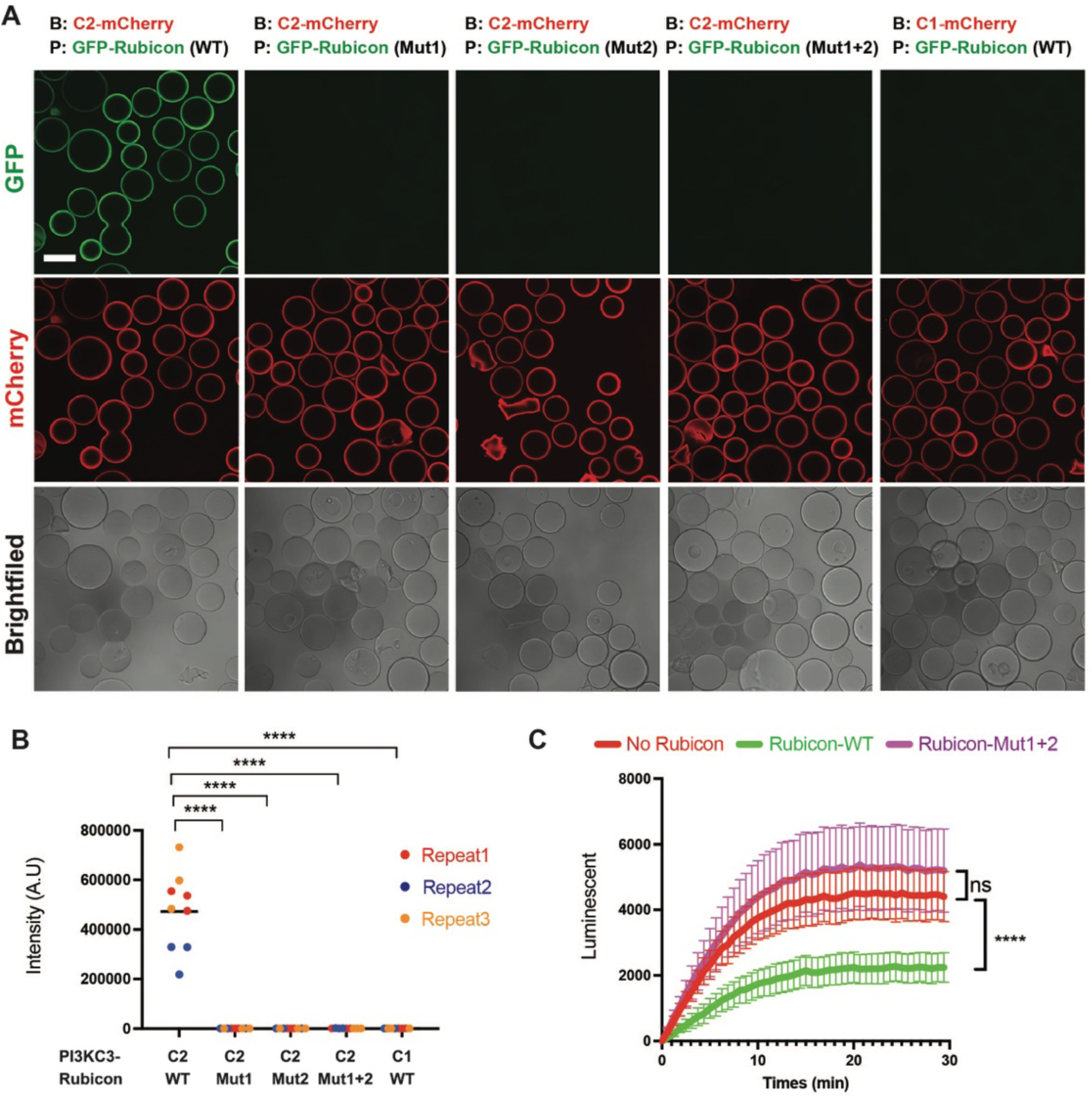
Interaction with Rubicon inhibits the activity of PI3KC3-C2 *in vitro*. (A) Bead-binding assay of PI3KC3 complexes and Rubicon fragments containing the helix α1 (478- 512). Strep beads coated with TwinStrep-Flag (TSF)- and mCherry-tagged PI3KC3 complexes were incubated with GFP-tagged WT Rubicon and its mutants. Mut1, E492K/S499K/E507K; Mut2, F496D/I503D. Scale bar: 100 μm. (B) Quantification of the Strep bead-binding assay. Statistical significance was determined by one-way ANOVA followed by Bonferroni’s multiple-comparison test. Each differently colored dot represents an independent biological replicate, and each biological replicate includes 3 technical replicates. The average fluorescence intensity of all beads in each image was calculated. A.U represents arbitrary units. ****P < 0.0001 and ns: not significant. Data are presented as mean ± s.d. (C) ATP consumption assay measuring PI3KC3-C2 activity on small unilamellar vesicles (SUVs) containing phosphatidylinositol (PI). Luminescent represents arbitrary units. ****P < 0.0001 and ns: not significant. Data are presented as mean ± s.d.

### PI3KC3-C2 subunit UVRAG defines the specific recognition of Rubicon by an indirect mechanism

The present Rubicon:PI3KC3-C2 complex structure demonstrates that the binding interface between the helix α1 of Rubicon and the BARA domain of BECN1 is sufficient for interaction. However, the PI3KC3-C2-specific subunit UVRAG does not directly participate in this binding interface. Thus, favorable direct interactions between UVRAG and Rubicon do not explain PI3KC3- C2 specificity. A precise comparison of PI3KC3-C2 with -C1 required us to determine the structure of PI3KC3-C1 in the absence of any other interactors that confound the comparison. The only available pre-existing structures of PI3KC3-C1 alone were at too low a resolution to support the detailed comparison needed (41, 42). For completeness, we also determined the structure of PI3KC3-C2 alone. We determined the unbound structures of the two complexes, each containing only the four core subunits (VPS34, VPS15, BECN1, and either ATG14 or UVRAG) at overall resolutions of 3.83 Å and 3.77 Å, respectively (SI Appendix Figs S4 and S5).

Although PI3KC3-C2 and C1 have the same overall architecture, PI3KC3-C2 adopts a slightly more extended conformation, resulting in a more L-shaped architecture compared to PI3KC3-C1 (SI Appendix Fig.S6A). Alignment of the VPS15^WD40^ scaffold domains revealed that the structural differences can be divided into two major regions. The most pronounced conformational changes occur within the base region, which contributes to the selective recruitment of Rab GTPases (SI Appendix Fig.S6B), as also seen in another recent report (37). In comparison, the adaptor arm underwent more subtle conformational rearrangements. This region consists of the VPS15^WD40^ scaffold, the mutually exclusive UVRAG^BARA2^ or ATG14^CTD^ domains, and the BECN1^BARA^ domain (SI Appendix Fig. S6C). Despite the structural similarity of the two BARA-like domains, UVRAG^BARA2^ contains an additional loop (residues 394-411) that engages the proximal coiled-coil (CC) side of the VPS15^WD40^ domain. In contrast, the C-terminal tail of the ATG14^CTD^ domain (residues 382-398) interacts with the distal CC side of the VPS15^WD40^ domain (SI Appendix Fig. S6C). These distinct binding modes result in a modest 3.3 Å displacement between the two BARA- like domains, which propagates to the distal BECN1^BARA^ domain, resulting in an 11.3° rotation and 8.5 Å translation between PI3KC3-C2 and C1 (SI Appendix Fig. S6D-E).

The Rubicon-binding surface on BECN1 remains essentially unchanged in the unbound PI3KC3- C2 structure (SI Appendix Fig. S2C), retaining the positively charged groove and the two hydrophobic pockets required for Rubicon binding. This observation indicates that the Rubicon- compatible conformation is an intrinsic structural feature of PI3KC3-C2 rather than one induced upon Rubicon association. In contrast, the corresponding surface undergoes substantial remodeling in the unbound PI3KC3-C1 structure, where the hydrophobic pockets are absent (Fig. 3A-B, SI Appendix Fig. S2D-E). One end of the groove, formed by the CC domain residues K265 and K266 together with the BARA domain residue N268 of BECN1, hereafter referred to as the ‘neck’ region, narrows from 9 Å in PI3KC3-C2 to 3 Å in PI3KC3-C1, thereby preventing accommodation of the Rubicon helix α1 (Fig. 3C-E). On the opposite face of the Rubicon-binding site, a hydrophobic patch on the BECN^BARA^ domain, composed of residues L264, V269, and A272, is stabilized by hydrophobic interactions with UVRAG residues F376 and F377 in PI3KC3-C2, or alternatively with ATG14 residues L210, I316, and I320 in PI3KC3-C1. Collectively, these observations indicate that the distinct architectures of the UVRAG^BARA2^ and ATG14^CTD^ domains allosterically determine the conformation of the BECN1 neck region, thereby conferring the specificity of Rubicon binding (Fig. 3F-H, Movie S1).

**Figure 3.**
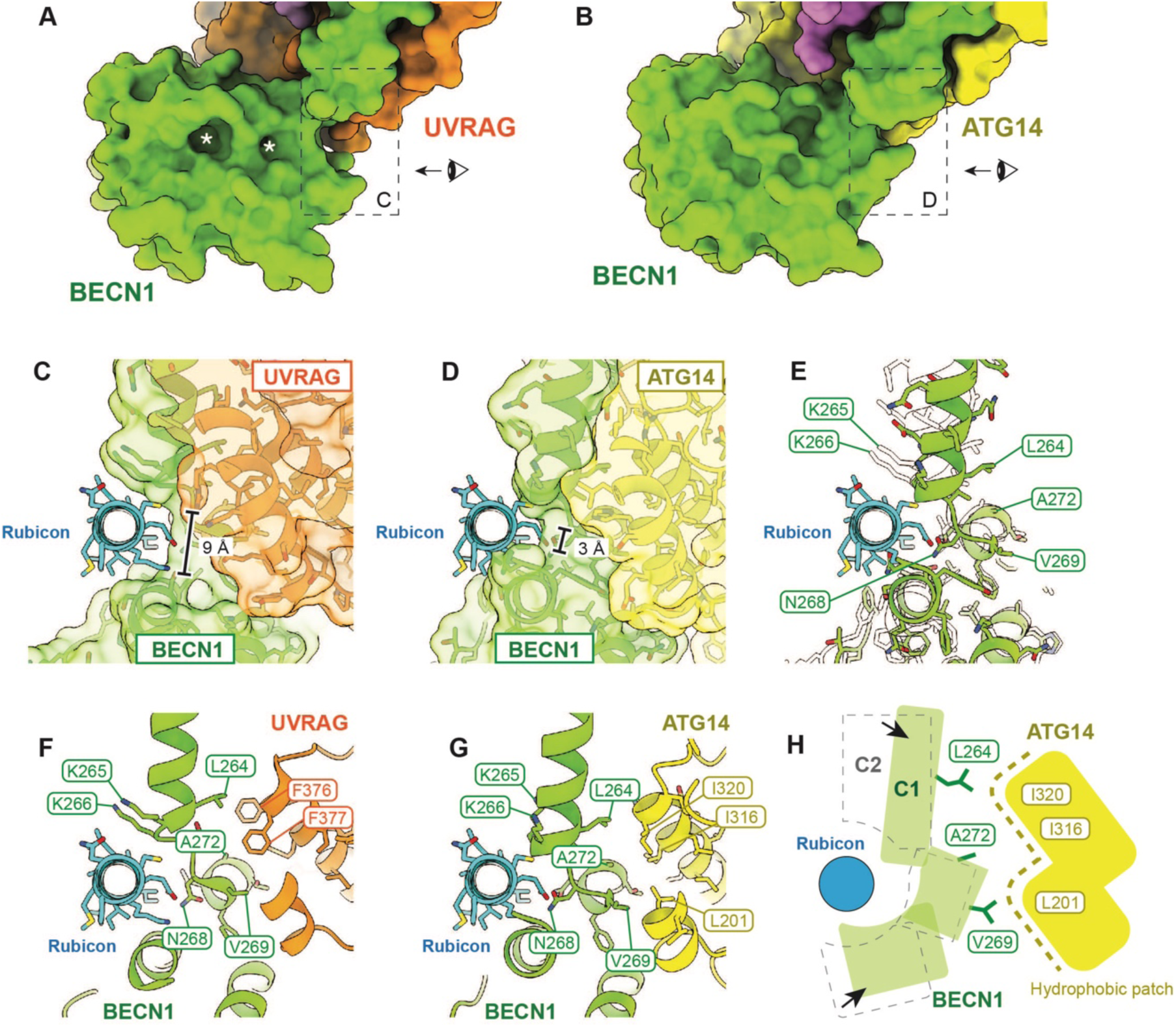
Structural basis for the PI3KC3-C2-specific interaction with Rubicon. (A) Close-up view of the Rubicon-binding site in the Rubicon complex. The Rubicon helix α1 is omitted to expose the BECN1^BARA^ surface. The two hydrophobic pockets are indicated by white asterisks. (B) Corresponding view of the Rubicon-binding site in the unbound PI3KC3-C1 structure. (C) Close-up view of the neck region in the Rubicon complex. The viewing region and orientation are indicated by the dashed rectangle in A. (D) Close-up view of the neck region in the unbound PI3KC3-C1 structure. The helix α1 of Rubicon is superimposed on the corresponding region of the unbound PI3KC3-C1 structure to illustrate the steric clash. (E) Superposition of the BECN1 subunit from the Rubicon complex (transparent) and the unbound PI3KC3-C1 structure (green). The Rubicon helix α1 is shown in cyan. Key BECN1 residues involved in Rubicon binding and formation of the hydrophobic patch are indicated. (F and G) Close-up views of the neck region in the Rubicon complex (F) and the unbound PI3KC3-C1 structure (G). The UVRAG^BARA2^ and ATG14^CTD^ domains located on the opposite face of the Rubicon-binding site are shown, with the key hydrophobic residues indicated. (H) Schematic model illustrating the allosteric regulation of the PI3KC3-C2- specific interaction with Rubicon. Incorporation of ATG14 (yellow) remodels the conformation of BECN1 from a Rubicon-compatible state (gray) to a Rubicon-repelling state (green). The hydrophobic patch and the corresponding residues are indicated. The critical conformational changes are indicated by arrows.

### Rubicon binding to PI3KC3-C2 suppresses mitophagy and lysosomal function

Using our previously established ARPE-19 mito-QC reporter cell line (43), we quantified mitophagy following Rubicon knockout (KO) and rescue with either wild-type (WT) Rubicon or the quintuple interaction-deficient mutant (Rubicon^Mut1+2^) described earlier (Fig. 4A). The mito-QC reporter consists of a tandem mCherry–GFP tag targeted to the outer mitochondrial membrane, allowing mitolysosomes to be identified as red-only puncta following lysosomal delivery, where GFP is quenched by the acidic environment while mCherry remains stable. Following treatment with deferiprone (DFP) to induce NIX/BNIP3-dependent mitophagy, or oligomycin/antimycin A (OA) to induce PINK1/Parkin-dependent mitophagy, cells expressing WT Rubicon exhibited significantly fewer mitolysosomes than Rubicon KO cells (Fig. 4B-C). In contrast, Rubicon^Mut1+2^ failed to suppress mitophagy, with mitolysosome numbers remaining comparable to those in Rubicon KO cells. Together, these findings demonstrate that disruption of the Rubicon–PI3KC3-C2 interaction abolishes the inhibitory effect of Rubicon on mitophagy, consistent with our biochemical analyses.

**Figure 4.**
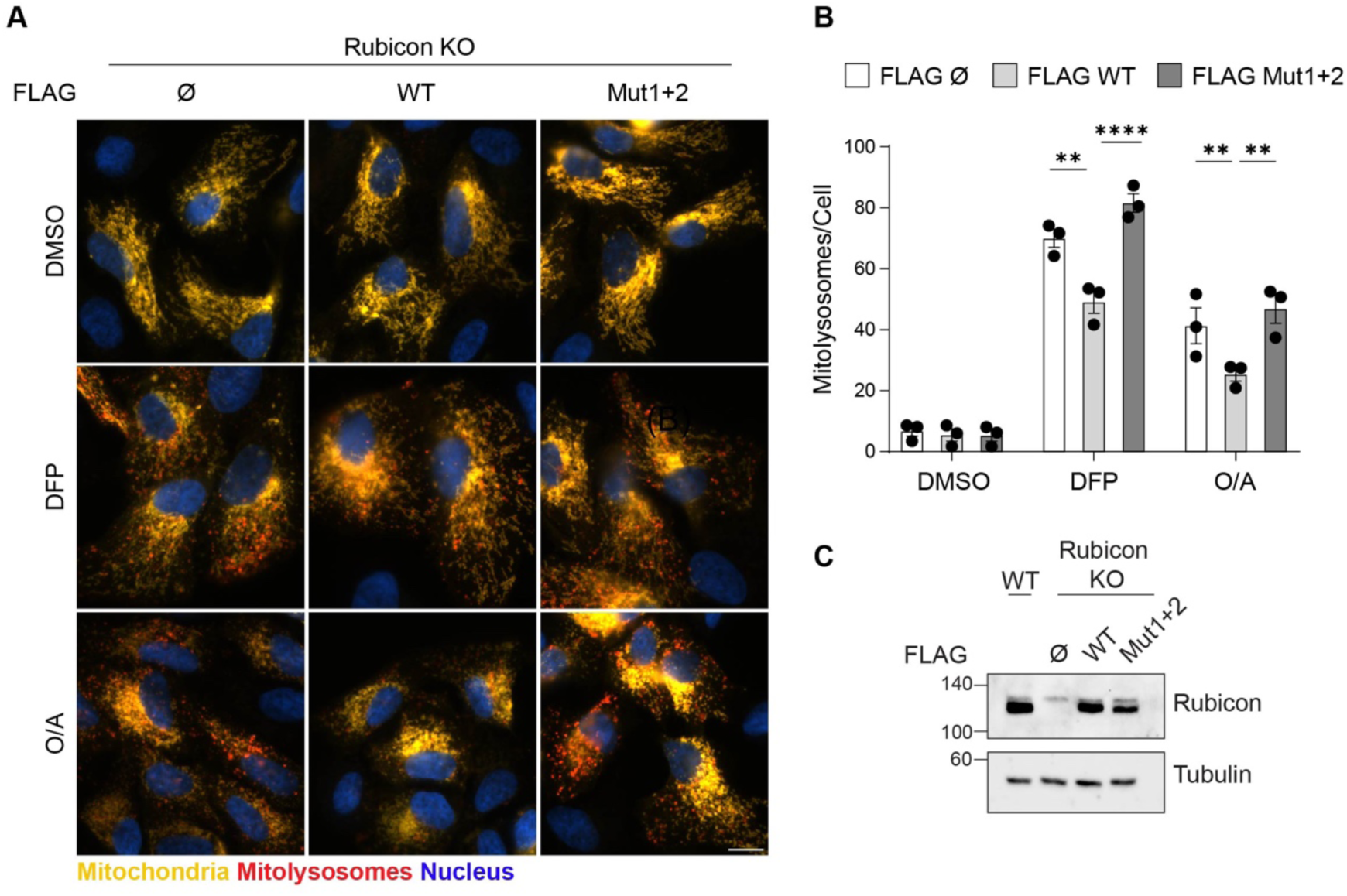
Interaction between Rubicon and PI3KC3-C2 suppresses mitophagy. (A) Representative wide-field images of Rubicon Knockout ARPE-19 mito-QC cells rescued with either FLAG-empty vector (Ø), WT, or Mut^1+2^ mutant Rubicon, treated with DFP (0.5mM) or a combination of oligomycin (1μM) and Antimycin (1μM) for 24 hours. Red puncta represent mitolysosomes while mitochondrial network appears yellow. Scale bar: 10 μm. (B) Quantification of the number of mitolysosomes (red puncta) per cell. Data represent mean ± SEM (n=3), significance calculated using ordinary two-way ANOVA and Tukey’s multiple comparisons test. Statistical significance is depicted as **p<0.01 and ****p<0.0001. (C) Representative immunoblots of indicated proteins from wildtype and Rubicon Knockout ARPE-19 cells rescued with either FLAG- empty, WT or M1+2 mutant Rubicon.

To further investigate the cellular basis of impaired mitophagy, we overexpressed either WT or Mut1 Rubicon in primary rat hippocampal neurons. Both WT and mutated Rubicons displayed comparable colocalization with LAMP1-positive organelles (Fig. 5A-C), indicating that disruption of the PI3KC3-C2 binding interface does not impair lysosomal recruitment, consistent with the previous finding that Rubicon is localized through its RH domain interaction with Rab7 (14, 29, 30).

**Figure 5.**
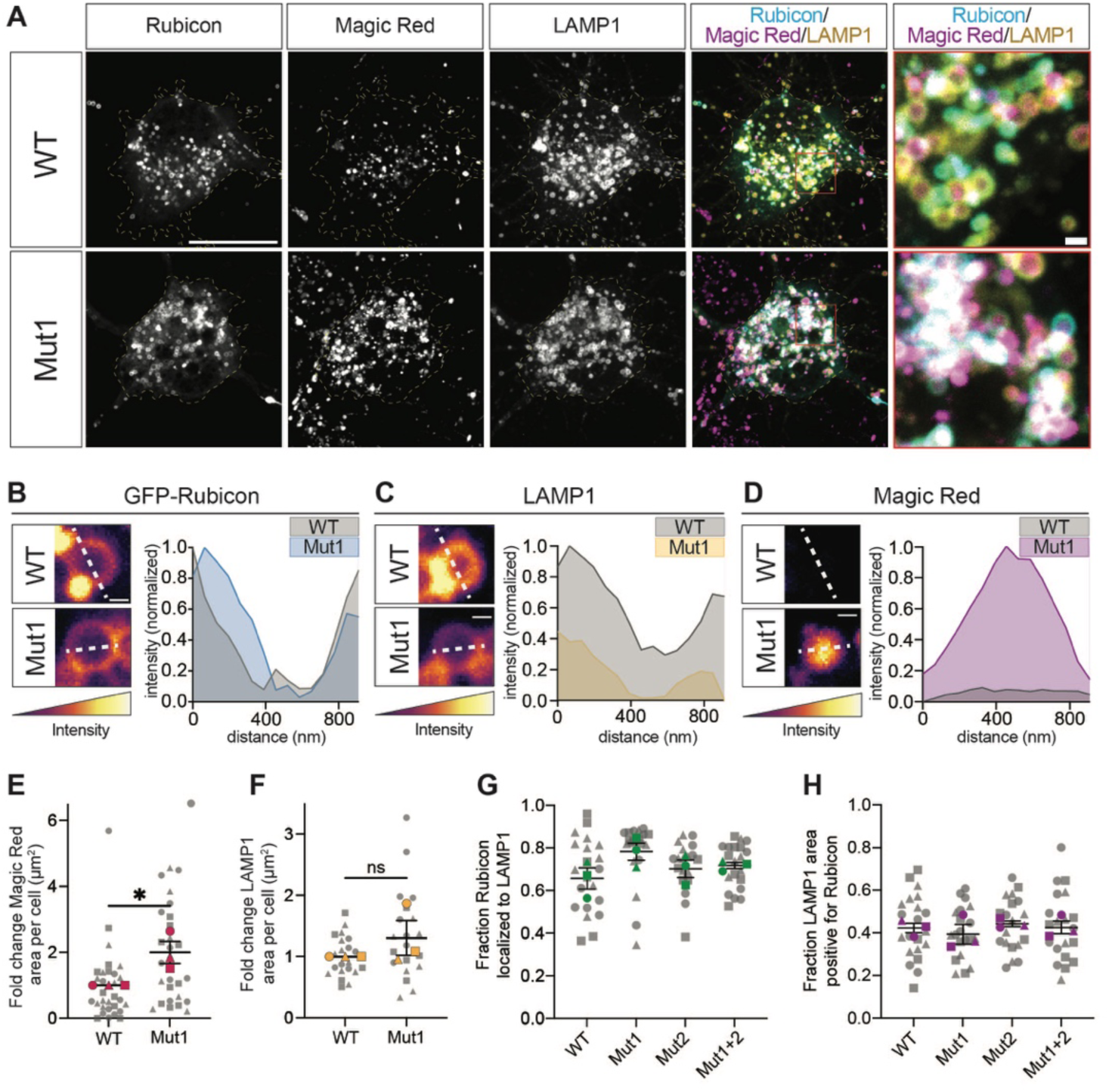
Interaction between Rubicon and PI3KC3-C2 suppresses lysosomal function. (A) Primary rat hippocampal neurons were transfected with either WT or Mut1 (E492K/S499K/E507K) GFP-Rubicon (cyan) together with Halo-LAMP1 (yellow) and stained with the cathepsin B activity reporter Magic Red (magenta). Scale bar: 20 μm; inset, 5 μm. (B–D) Line- scan analysis of normalized fluorescence intensity across vesicles positive for GFP-Rubicon and Halo-LAMP1 in neurons expressing WT or Mut1 Rubicon. Scale bar, 500 nm. (E) Magic Red- positive area per neuron, normalized to the corresponding WT condition. (F) LAMP1-positive area per neuron, normalized to the corresponding WT condition. Gray symbols represent individual technical replicates, whereas colored symbols represent independent biological replicates (*n* = 3). Statistical significance was determined using two-tailed unpaired *t*-tests on biological replicate values. *P* < 0.05. (G) Manders coefficient analysis representing the fraction of somal Rubicon localized to LAMP1-positive organelles. (H) Manders coefficient analysis representing the fraction of somal LAMP1 area that is positive for Rubicon. Graphs depict individual technical replicates shown in gray and biological replicates (n=3) shown in color. Significance relative to WT control was determined by one-way ANOVA with multiple comparisons performed on independent biological replicate values. Error shown as mean +/- SEM.

We next assessed lysosomal proteolytic activity using the fluorogenic cathepsin B substrate Magic Red. Neurons expressing Rubicon^Mut1^ exhibited a significant increase in active lysosomes compared with those expressing WT Rubicon (Fig. 5A,D-E), whereas the total lysosomal area remained unchanged (Fig. 5F). The lysosomal recruitment of Rubicon and its mutants was assessed by quantifying the fraction of LAMP1-positive Rubicon (Fig. 5G) and the fraction of Rubicon-positive LAMP1 (Fig. 5H). No significant changes were observed between the WT and Mut1, as well as the Mut2 and the quintuple mutant Mut1+2. This is consistent with the main determinant of lysosomal localization mapping to the Rab7-binding RH domain, remote from the PIKBD and the sites of mutation. These findings indicate that Rubicon mutants are recruited to lysosomes as efficiently as WT Rubicon but are substantially less effective at suppressing lysosomal function. Together, our data establish that direct interaction between Rubicon and PI3KC3-C2 is required to inhibit PI3KC3-C2 activity, thereby suppressing mitophagy and lysosomal function.

## Discussion

Here, we set out to understand why Rubicon only binds to PI3KC3-C2. We found that Rubicon does not directly contact the C2-unique UVRAG subunit. Nor is there direct steric overlap between the C1-unique subunit ATG14 and Rubicon. Instead, the mutually exclusive subunits UVRAG and ATG14 allosterically remodel the conformation of the shared BECN1^BARA^ domain. UVRAG promotes a BECN1 conformation capable of binding Rubicon PIKBD helix α1, while ATG14 does the opposite. This finding uncovers an unexpected mechanism by which complex-specific subunits confer interaction specificity through conformational regulation of a common scaffold rather than by directly participating in ligand binding.

The PIKBD domain of Rubicon was previously identified as the region responsible for binding PI3KC3-C2 (23), but the atomic details of this interaction were not defined well enough to surgically excise this biochemical activity and thereby probe its role in cell physiology. The high-resolution cryo-EM structure presented here enabled us to carry out structure-guided mutagenesis and so selectively disrupt the Rubicon-PI3KC3-C2 interface. Disruption rescued PI3KC3-C2 enzyme activity against liposomes *in vitro*, rescued mitophagy to levels comparable to those observed in Rubicon KO cells (Fig. 4), and enhanced lysosomal proteolytic activity without altering lysosomal abundance (Fig. 5). These findings indicate that the negative regulatory function of Rubicon in both mitophagy and lysosomal function can be accounted for by its ability to interact with and inhibit PI3KC3-C2.

PI3KC3-C2 is best known for its role as a driver of endolysosomal maturation; thus, it makes sense that Rubicon-mediated inhibition of PI3KC3-C2 has a negative effect on lysosomal function. The findings here add further detail to the observation that Rubicon suppresses lysosomal activity in neurons (14) by showing that this effect is mediated by C2 inhibition. These findings are consistent with the finding that Rubicon acts to inhibit autophagy mainly at the autophagosome-lysosome fusion step (14). The most parsimonious interpretation of the effect of Rubicon on mitophagic flux is that decreased mitochondrial turnover is a downstream consequence of reduced ELN maturation. A full understanding of the interplay of Rubicon’s negative effects on both autophagy and endolysosomes will require further work. The dual role of Rubicon in both processes suggests that antagonizing Rubicon could have broad benefits in increasing the degradative capacity of cells.

Our findings also provide a framework for understanding how Rubicon suppresses lysosomal function. Previous studies have established that PI(3)P is essential for lysosomal fusion through the recruitment of the HOPS (Homotypic fusion and vacuole protein sorting) complex (44) and SNARE (Soluble N-ethylmaleimide-sensitive factor Attachment Protein Receptors) proteins (45). PI(3)P also serves as the precursor for the synthesis of PI(3,5)P2 by PIKfyve (46, 47), which is central to lysosomal functions including fusion. We therefore propose that Rubicon is recruited to lysosomes via Rab7 (12, 27) while its PIKBD captures adjacent PI3KC3-C2 complexes and restricts their productive association with membranes. This inhibition is predicted to suppress local PI(3)P production, thereby impairing recruitment of membrane fusion machinery and reducing lysosome-autophagosome fusion. In this scenario, reduction in mitophagy and other types of autophagy are downstream consequences of reduced lysosome-autophagosome fusion, as well as reduced ELN activity.

Rubicon has been implicated in a range of biology beyond the ELN and autophagy. Rubicon also regulates exosome biogenesis (48), LC3-associated phagocytosis (LAP) (49, 50), and modulates neuroimmune responses in traumatic brain injury (51). A shortened form of Rubicon is reported to positively regulate autophagy in B cells (52). The determinants of these processes within the Rubicon molecule have yet to be mapped in detail. While Rubicon KO at the whole animal level appears to have few consequences that would be seemingly therapeutically undesirable (13), it will nevertheless be important to gain a deeper understanding of the putative non-autophagy and non-lysosomal consequences of Rubicon inhibition. To the extent there are, it will be important to determine if these roles are connected to PI3KC3-C2. The Rubicon mutants described here should be valuable to uncover whether PI3KC3-C2 interaction mediates these processes.

In summary, our study establishes the atomic basis for the selective interaction between Rubicon and PI3KC3-C2 and identifies differential allosteric remodeling of the shared core subunit BECN1 as the mechanism underlying complex II specificity. By combining structural, biochemical, and cellular analyses, we demonstrated that this interaction is essential for Rubicon-mediated inhibition of lysosomal function and autophagy. These findings provide a mechanistic framework for understanding how PI3KC3-C2 is selectively regulated and suggest that disruption of the Rubicon:PI3KC3-C2 interface may represent a more selective therapeutic strategy for autophagy and endolysosome enhancement, as opposed to the wholesale inhibition or degradation of Rubicon.

## Materials and Methods

### Plasmid construction

Plasmids pCAG-VPS34 (Addgene#203564), pCAG-VPS15-TSF (Addgene #203563), pCAG- BECN1 (Addgene#203568), and pCAG-mCherry-ATG14(#203567) were from Addgene. pGST2- GFP-Rubicon-PIKBD-WT, pGST2-GFP-Rubicon-PIKBD-Mut1 (E492K/S499K/E507K), pGST2-GFP-Rubicon-PIKBD-Mut2 (F496D/I503D) and pGST2-GFP-Rubicon-PIKBD- Mut1+2 were synthesized via Integrated DNA Technologies and subcloned into pGST2-GFP vector using recombination. To get the linear fragment of pGST2-GFP, the following primers were used: pGST2- GFP-F: GTCGACGAGCTCACTAGTCGC, pGST2-GFP-R: CGCTTCGTCGCTACCGGA. To generate pCAG-mcherry-UVRAG and pCAG-UVRAG, human UVRAG and mcherry-UVRAG were synthesized via Integrated DNA Technologies and subcloned into pCAG vector using CalI and XhoI. pCAG-BECN1-(GSS)4-Rubicon (472-672) was from our previous study (23). pGST2-GFP-Rubicon-PIKBD-WT, pGST2-GFP-Rubicon-PIKBD-Mut1 (E492K/S499K/E507K), pGST2-GFP-Rubicon-PIKBD-Mut2 (F496D/I503D) and pGST2-GFP-Rubicon-PIKBD-Mut1+2, pCAG-mcherry- UVRAG and pCAG-UVRAG were confirmed by sequencing and deposited to Addgene. pBABE mito-QC retroviral system (#DU40799) was used as described previously (53). pBabeD hygro FLAG Rubicon Wild-type (#DU78825), pBabeD hygro FLAG Rubicon E492K F496D S499K I503D E507K (Mut1+2, #DU83454) retroviral constructs were obtained from MRC-PPU reagents and Services. Details are shown in the key resource table.

### Cell culture for protein expression

HEK293F GnTi cells were maintained in FreeStyle™ 293 Expression Medium (Gibco) supplemented with 1% FBS and 1x anti-anti. Cells were treated under the indicated conditions and maintained at 37℃ in 5% CO2.

ARPE-19 cell lines were grown at 37°C with 5% CO2 in a humidified incubator. Wild-type and Rubicon KO ARPE-19 cells were passaged and maintained in DMEM: F12 media supplemented with 10% Fetal bovine serum, 2mM L-glutamine, 100 U/ml Penicillin, and 0.1 mg/ml Streptomycin. Stable expression of the protein of interest in ARPE-19 was generated using Retroviral transduction. Viral particles were produced in 70% confluent HEK293FT by transfecting 1 µg of pBABE vector expressing the respective genes, 0.66 µg of Gag-Pol, and 0.34 µg of VSVG using Lipofectamine 2000 (Thermo Scientific) according to the manufacturer’s protocol. The cell supernatant was harvested after 48 hours of transfection and filtered through a 0.45 μm filter to remove floating cells. The filtered supernatant was used to transduce ARPE-19 cells at 70% confluency with polybrene (10 μg/ml). After 48 hours of transduction, the cells were selected with either puromycin (2 μg/ml) or hygromycin (100 μg/ml), and the stable pool of cells was used for the experiment.

### Generation of Rubicon KO CRISPR cell line

Paired CRISPR guides targeting exon 5, cloned in the pX335 Cas9 vector and pBABED_P_U6 vector, were used to generate RUBICON knock-out ARPE-19 cells. Sense Guide: GTGACATGAGCGTCACTTAGC and Anti-Sense: GCTGCAGTGCCTGGAAGCAG were cloned into the respective guides. 5 μg of each endotoxin-free guide was electroporated into ARPE-19 cells using the Super Electroporator NEPA21 system at 175V for 5ms. Following electroporation, cells were incubated overnight in antibiotic-free media under standard culture conditions. After 24 hours, positive cells were selected with 2 µg/ml of puromycin for 24 hours. After recovery, the CRISPR pool was used to check KO efficiency, and single-cell clones were generated by serial dilution and screened using immunoblotting for Rubicon.

To confirm Rubicon knockout, genomic DNA was extracted using the Qiagen DNeasy kit following the manufacturer’s instructions. The CRISPR cleavage site was PCR amplified using Platinum™ SuperFi II PCR Master Mix (Thermo Scientific) according to the manufacturer’s protocol and checked in 1% Agarose gel. The PCR product was then purified using Agencourt AMPure XP (Beckman Coulter) magnetic beads. Samples were then analyzed by mi-Sequencing, which revealed a 5-bp insertion at the 109th codon in one allele and a 16-bp insertion in the second allele, causing a frameshift mutation.

### Protein expression and purification

For the cryo-EM sample of PI3KC3-C2, pCAG-VPS34, pCAG-VPS15-TSF, pCAG-BECN1, and pCAG-UVRAG were co-transfected into HEK293 GnTi^-^ cells at a 1:1:1:1 mass ratio using the polyethylenimine (PEI) (Polysciences) transfection system. For the cryo-EM samples of PI3KC3- C2:Rubicon complex and PI3KC3-C1, the pCAG-BECN1 was substituted with pCAG-BECN1- (GSS)_4_-Rubicon(472-672), and the pCAG-UVRAG was substituted with pCAG-ATG14, respectively. For the beads binding assay or GUV assay, pCAG-VPS34, pCAG-VPS15-TSF, pCAG-BECN1, and pCAG-mcherry-UVRAG (PI3KC3-C2 complex) or pCAG-mcherry-ATG14 (PI3KC3-C1 complex) were used.

1 L HEK293 GnTi^-^ cells were transfected at a concentration of 2 × 10^6^/ml and harvested after 48 hours. The cells were pelleted by centrifugation at 1,500 × g, washed with phosphate-buffered saline (PBS), spun again at 500 × g for 10 min in a tabletop centrifuge, flash-frozen, and stored at −80 °C for later use. Cell pellets were thawed at room temperature and resuspended in purification buffer containing 25 mM 4-(2-Hydroxyethyl)piperazine-1-ethanesulfonic acid (HEPES) pH 7.5, 200 mM NaCl, 2 mM MgCl_2_, and 10 mM tris(2-carboxyethyl)phosphine (TCEP). An EDTA-Free Protease inhibitor tablet (Thermo Scientific) was added, and the gently resuspended pellet was transferred to a Pyrex Dounce homogenizer. The cells were Dounce homogenized 20 times. The homogenate was transferred to a new tube, and glycerol and Triton X-100 were added to the cells at final concentrations of 10% and 1%, respectively, gently mixed, and then left to rock at 4 °C for 1 hour. Following detergent lysis, the cells were pelleted by centrifugation (40,000 × g for 60 minutes at 4 °C). The supernatant was collected and passed through a 0.22 μm filter (Avantor Sciences) to remove invisible aggregates, then mixed with 2 mL pre-washed Strep-Tactin Sepharose resin (IBA Lifesciences) at 4°C overnight. The following day, the resin was washed with 20 ml of purification buffer until the post-column flow was protein-free, as confirmed by SDS-PAGE. The bound protein was then eluted with 10 ml of purification buffer containing 10 mM desthiobiotin. Samples were concentrated and then loaded onto a Superose 6 Increase 10/300 GL column (Cytiva) equilibrated in purification buffer. The concentration of the eluted protein samples was measured by Nanodrop (Thermo Scientific). The fresh sample was used for cryo-EM preparation or aliquoted into 25 μl fractions, flash-frozen in liquid nitrogen, and stored at −80 °C until use in other assays.

GST-GFP-Rubicon-PIKBD-WT, GST-GFP-Rubicon-PIKBD-Mut1 (E492K/S499K/E507K), GST-GFP-Rubicon-PIKBD-Mut2 (F496D/I503D) and GST-GFP-Rubicon-PIKBD-Mut1+2 were expressed and purified from *E.coli*. Plasmids were transformed into BL21 competent cells, and a single clone was picked and grown in 1 L LB broth. Cell were harvested by centrifugation at 4,000 × g, washed with phosphate-buffered saline (PBS), spun again at 4,000 × g for 10 min in a tabletop centrifuge, flash-frozen, and stored at −80 °C for later use. Cell pellets were thawed at room temperature and resuspended in purification buffer containing 50 mM 4-(2- Hydroxyethyl)piperazine-1-ethanesulfonic acid (HEPES) pH 7.5, 200 mM NaCl, 2 mM MgCl2,, 2 mM tris(2-carboxyethyl)phosphine (TCEP) and 5% glycerol. An EDTA-Free Protease inhibitor tablet (Thermo Scientific) was added. After sonication (40% power, 2s on, 2s off) for 10 min, the lysates were pelleted by centrifugation (40,000 × g for 60 minutes at 4 °C). The supernatant was collected and passed through a 0.22 μm filter (Avantor Sciences) to remove invisible aggregates, then mixed with 2 mL pre-washed Glutathione Sepharose 4B (GE Healthcare) at 4°C for 3h. Then, the resin was washed with 150 ml of purification buffer until the post-column flow was protein-free, as confirmed by SDS-PAGE. The bound protein was then eluted with 20 mL of purification buffer containing 50 mM glutathione (pH 7.5). Samples were concentrated and then loaded onto a S200 10/300 column (Cytiva). The running buffer is 50 mM HEPES, 200 mM NaCl, 2 mM MgCl_2_, 1 mM TCEP. The concentration of the eluted protein samples was measured using a NanoDrop (Thermo Scientific). The fresh sample was aliquoted into 100 μl fractions, flash-frozen in liquid nitrogen, and stored at −80 °C until used for other assays.

### Sample vitrification and cryo-EM data acquisition

For cryo-EM sample preparation, 3 μl of protein solution at a concentration of 0.25-0.45 mg/ml was applied onto freshly glow-discharged grids (QUANTIFOIL R1.2/1.3 or R2/1 mesh 300, Electron Microscopy Sciences) in the PELCO easiGlow system (Ted Pella) at 25 mA current for 30 sec. Sample vitrification was performed with a Vitrobot cryo-plunger (Thermo Fisher Scientific) at 100% humidity, 4 °C, a 3 sec wait time, and a blot force of −10. 0.05% (w/v) n-Octyl-β-D-glucopyranoside (Anatrace) was added to the sample solution as a surfactant prior to vitrification.

The datasets of the unbound PI3KC3-C2 and Rubicon:PI3KC3-C2 complexes were recorded on a 300 kV Titan Krios microscope equipped with an X-FEG and energy filter set to 20 eV. The data were automatically collected with SerialE(54) on a K3 Summit direct electron detector (Gatan) at an 81,000x magnification, with super-resolution pixel sizes of 0.525 Å and 0.470 Å, respectively, and a defocus range of −0.8 to −2.0 micrometers. 50-frame image stacks were collected to a final cumulative dose of ∼50 e/Å^2^. The unbound PI3KC3-C1 dataset was recorded on a 200 kV Talos Arctica microscope equipped with an X-FEG. The data were automatically collected with SerialEM (54) on a K3 Summit direct electron detector at a magnification of 36,000x, with a super-resolution pixel size of 0.558 Å and a defocus range of −0.8 to −2.0 micrometers. 50-frame image stacks were collected to a final cumulative dose of ∼50 e/Å^2^. Other details of the dataset collection are summarized in SI Appendix S1.

### Image processing and 3D reconstruction

The datasets were processed using the cryoSPARC (55) workflow. In brief, the super-resolution video stacks were motion-corrected and binned 2x by Fourier cropping using Patch Motion Correction. Contrast transfer function determination was performed using Patch CTF Estimation, followed by manual removal of outlier micrographs based on the estimated defocus and resolution values. Single particles were automatically picked by Topaz (56) based on a manually trained model, extracted using a window size that is 1.5-2 times larger than the target particle, and further binned to 2-4x to facilitate subsequent processing. Two-dimensional (2D) classification was then used to remove obvious junk particles before subsequent classification. The initial maps were obtained by using *ab initio* reconstruction. Further classification was done at the 3D level by multiple rounds of heterogeneous refinement until a clean substack was obtained. The particles were re-extracted from the micrographs using the refined coordinates at the original 2× pixel bin size and were then used for homogeneous refinement over multiple rounds until the final resolution converged. To further improve the quality of the map, local refinement was applied using masks generated by the Volume Tools in UCSF ChimeraX (57). Each local map was aligned to the consensus map and composed using the ‘vop maximum’ command in UCSF ChimeraX. The composed maps were then used to build the model. The details of data processing are summarized in SI Appendix Table S1.

### Model building, validation, and visualization

The *in silico* models of unbound PI3KC3-C2 and PI3KC3-C1 were generated using AlphaFold2 predictions(58). The resolution of the observed maps enabled amino acid sequence assignment. The primary and secondary structures of the predicted models agree well with the EM maps. A flexible model was fitted using the real-time molecular dynamics simulation-based program ISOLDE(59), implemented in the visualization software UCSF ChimeraX, followed by iterative refinement with the model editing software Coot(60) and real-space refinement in Phenix(61). The electrostatic potentials on molecular surfaces were calculated using APBS(62), and the hydrophobic properties of the surfaces were calculated using the ‘mlp’ command in UCSF ChimeraX.

### Microscopy-based bead-binding assay

A mixture of 1 µM purified GST-tagged protein and 500 nM purified fluorescent protein in a total volume of 70 µl was incubated with 9 µl of preblocked glutathione Sepharose beads (Cytiva) in a reaction buffer containing 25 mM HEPES at pH 7.5, 150 mM NaCl, 1 mM MgCl2 and 1 mM TCEP. After incubation at room temperature for 30 min, samples were mixed with an additional 100 µl of reaction buffer and then transferred to the observation chamber for imaging. Images were acquired on a Nikon A1 confocal microscope with a Nikon Plan APO VC ×20 0.75 numerical aperture ultraviolet microscope objective. Three biological replicates were performed for each experimental condition.

### SUV production and lipid kinase assay

Liposomes were prepared by first drying overnight and rehydrating 0.43 mg of lipids (40% DOPC, 20%DOPE, 20% DOPS, and 20% Liver PI) with 860 μL of buffer containing 25 mM HEPES (pH 7.5), 200 mM NaCl, 2mM Mgcl_2_ and 2mM TECP on ice, resulting in a final concentration of 0.6 mM total lipid (0.5 mg/mL). The lipid suspension was vortexed for 5 minutes and subjected to 10 freeze-thaw cycles. The rehydrated lipids were then extruded 21 times through a 50-nm polycarbonate membrane using a mini extruder, yielding 50-nm SUVs.

The lipid kinase assay was carried out using ADP-Glo Kinase Assay (Promega). Freshly purified PI3KC3-C2 and Rubicon fragment were pre-incubated with SUVs in the reaction buffer (50 mM HEPES pH 7.5, 200 mM NaCl, 2 mM MnCl2, 2mM TCEP). The reaction was initiated by adding 2ul ATP (250 mM), and incubated at room temperature for 30 min. An ATP-depletion reagent was added to terminate the lipid kinase reaction for 1h, deplete the remaining ATP, leaving only ADP. Then a kinase detection reagent was added to convert ADP to ATP, which is used in a coupled luciferase reaction. The luminescent output was measured with an INFINITE M PLEX (TECAN).

### Microscopy-based mito-QC assay in cell lines

Rubicon KO and rescue cells stably expressing the mito-QC sensor were seeded on a coverslip and incubated overnight in standard culture conditions. After 24 hours, respective drug treatments were performed for 24 has mentioned above. At end of treatments, the cells were washed with PBS (three times) and fixed for 10 min at room temperature using 3.7% paraformaldehyde (PFA) in 200 mM HEPES buffer (pH 7.0). To quench residual PFA, the cells were then washed and incubated in DMEM supplemented with 10 mM HEPES (pH 7.0) and 0.02% sodium azide (NaN₃) for 15 min at room temperature. Coverslips were then washed twice with PBS, mounted using ProLong™ Glass Antifade Mountant (Invitrogen) and imaged with a Nikon Eclipse Ti2 widefield microscope equipped with a 60X objective. The number of mitolysosomes indicated by red puncta was quantified using the semi-automated mito-QC counter plugin in ImageJ, as previously described (43).

### Immunoblotting

Cells were lysed in NP-40 lysis buffer (50 mM HEPES (pH 7.4), 150 mM NaCl, 1 mM EDTA, 10% glycerol, and 1% NP-40) supplemented with phosphatase inhibitors (1.15 mM sodium molybdate, 4 mM sodium tartrate dihydrate, 10 mM β-glycerophosphoric acid disodium salt pentahydrate, 1 mM sodium fluoride, and 1 mM activated sodium orthovanadate) and a protease inhibitor cocktail (Roche). Total protein concentrations were measured using the Pierce BCA Protein Assay Kit (Thermo Scientific). Equal amounts of protein were separated by SDS-PAGE on 4–15% Bis-Tris gradient gels at 120 V for approximately 90 min using 1× Bis-Tris running buffer. Proteins were subsequently transferred onto 0.45 µm Amersham™ Protran® nitrocellulose membranes (Cytiva). Membranes were blocked in 5% skimmed milk in TBST and incubated with the respective primary antibodies overnight at 4°C. After incubation membranes were washed with TBST three times, membranes were then incubated with HRP-conjugated secondary antibodies for 1 h at room temperature. Next membranes were again washed in TBST thrice and imaged using Clarity Western ECL Substrate (Bio-Rad) in Bio-Rad ChemiDoc imaging system according to the manufacturer’s instructions.

### Neuronal culture and transfection

Hippocampal neurons isolated from embryonic day 18 Sprague-Dawley rats were obtained from the Neurons R Us Culture Service Center (RRID: SCR_022421) at the University of Pennsylvania. Neurons (200k-250k/dish) were plated on glass-bottomed imaging dishes precoated with 0.5mg/ml poly-L-lysine (Sigma-Aldrich). Cells were initially plated in MEM (Gibco) containing 10% horse serum, 33 mM D-glucose, 1 mM sodium pyruvate); once adherent, this media was replaced with maintenance media consisting of neurobasal (Gibco) with 33 mM D-glucose (Sigma-Aldrich), 2 mM GlutaMAX (Thermo Fisher Scientific), 100 U/ml penicillin and 100 mg/ml streptomycin (Gibco), and 2% B-27 5x supplement (Thermo Fisher Scientific). Neurons were maintained at 37°C in 5% CO_2_ and AraC (1 µM) was added 24 hours after plating.

Between days in vitro (DIV) 6-9, neurons were transfected for 24-48 hours using Lipofectamine 2000 per the manufacturer’s protocol. Cells were transfected with 0.75 μg of the indicated GFP-Rubicon plasmid, in addition to 0.25 μg LAMP1-Halo. Thirty minutes prior to imaging, neurons were treated with Magic Red Cathepsin B Assay Kit (BioRad) as previously described(14). Hippocampal neurons were imaged in Hibernate E medium (BrainBits) supplemented with 2% B-27 and 33mM D-glucose containing 20nM Janelia Fluor 646-Halo ligand (Promega).

### Spinning disk confocal imaging and quantitative image analysis

Neurons were imaged using an Orbital-200 CSU spinning disk confocal mounted on a Nikon Eclipse Ti stand operated by VisiView Image Acquisition software, using an apochromatic 100x 1.49 NA oil immersion objective and a Hamamatsu CMOS ORCA-Fusion (C11440-20UP) camera. Images were acquired at 200nm intervals throughout the volume of each neuron, and image analysis was carried out using Fiji Is Just ImageJ(63) on max projections of each channel. Somas were manually traced, and only the content within the soma of each neuron was quantified. To determine cellular area positive for Magic Red staining, control images of non-transfected neurons were used to determine a suitable threshold above which Magic Red staining appeared accurate and luminal. Magic Red images were then uniformly converted to binary, with everything above that intensity threshold treated as a Magic Red-positive pixel, and the area of Magic Red-positive pixels per cell was analyzed. LAMP1 and Rubicon images underwent background subtraction with a rolling-ball radius of 50, and binary images for each were manually and blindly generated using the Otsu threshold prediction for each cell. The LAMP1 area per cell was determined from these binaries. All binary images were passed through the despeckle function in FIJI. Manders coefficients for Rubicon localization to LAMP1 in the soma were determined by using these binaries to calculate (area positive for both LAMP1 and Rubicon)/(total area positive for Rubicon). Manders coefficients for LAMP1 area positive for Rubicon were determined by using these binaries to calculate (area positive for both LAMP1 and Rubicon)/(total area positive for LAMP1).

To create normalized line scans for visualization, a 910nm trace was created across a single representative lysosome from each condition, and the intensity for each channel as a function of distance was recorded. All pixel intensities for each condition were normalized to the highest intensity value recorded in that channel in either condition. The lowest normalized value was then uniformly subtracted, and all values were again divided by the highest normalized intensity value present for each channel.

### Statistical Analysis for the primary neuron assays

Neurons obtained through separate dissections and transfected separately were considered independent biological replicates. Statistical analyses were performed on the averages from all images within each biological replicate.

### Data accessibility

The cryo-EM maps have been deposited in the Electron Microscopy Data Bank (EMDB) under the following accession codes: Rubicon:PI3KC3-C2 complex: PDB: 9ZPD, EMD-74527, EMD-74522, EMD-74520, EMD-74521; PI3KC3-C2 unbound form: PDB: 9ZPC, EMD-74526, EMD-74511, EMD-74509, EMD-74510; PI3KC3-C1 unbound form structure: PDB:13BV, EMD-76952, EMD- 76934, EMD-76932, EMD-76933. Protocols are available on https://www.protocols.io/. Plasmids developed for this study have been deposited at https://www.addgene.org/. Original gel scans, ADP-Glo data, and light microscope images shown in this study have been deposited at Zenodo. The details of data accessibility are summarized in the SI Appendix Table S2.

## Supporting information

Key resources table

## Acknowledgments

We thank all members of the Hurley lab and Aligning Science Across Parkinson’s (ASAP) team Mito911 for advice and discussions. We thank K. Sharma for his support of the cryo-EM facility. This research was funded by Aligning Science Across Parkinson’s (ASAP-000350) through the Michael J. Fox Foundation for Parkinson’s Research (to E.L.F.H., I.G.G., and J.H.H.) and the National Institutes of Health (R01 NS134598 to J.H.H.).

## Author Contributions

Conceptualization, J.H.H.; Methodology, M.C., A.B., Y.D., J.F.R., Q.N., A.J., I.J.A.; Investigation, M.C., A.B., Y.D., J.F.R.; Visualization, M.C., A.B., Y.D., J.F.R.; Supervision, E.L.F.H., I.G.G., J.H.H. Writing - original draft, M.C. and J.H.H. Writing - review and editing, all authors.

## Supporting Information for

**Fig. S1.**
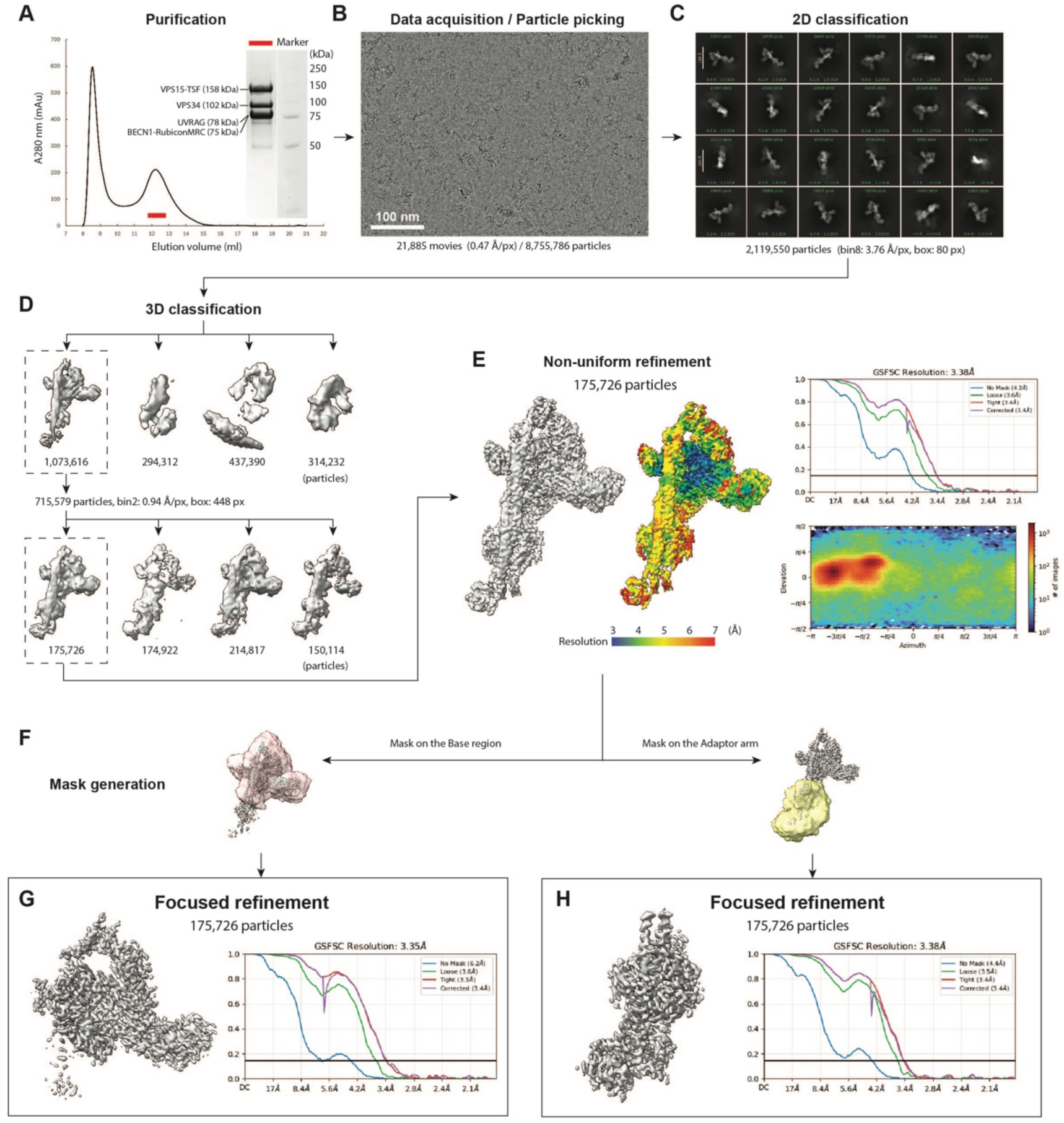
Cryo-EM sample preparation, image acquisition, and data processing of the Rubicon:PI3KC3-C2 complex. (A) Size-exclusion chromatography (SEC) profile of the Rubicon:PI3KC3-C2 complex. The inset shows an SDS–PAGE analysis of the peak fraction (red bar). (B) Representative cryo-EM micrograph. (C) Representative 2D class averages. (D) Results of the initial and final rounds of 3D classification. (E) Consensus refinement of the final particle stack, with the Fourier shell correlation (FSC) curve and local resolution map. (F) Mask generation for focused refinement. (G and H) Results of focused refinement, with the corresponding cryo-EM maps and FSC curves.

**Fig. S2.**
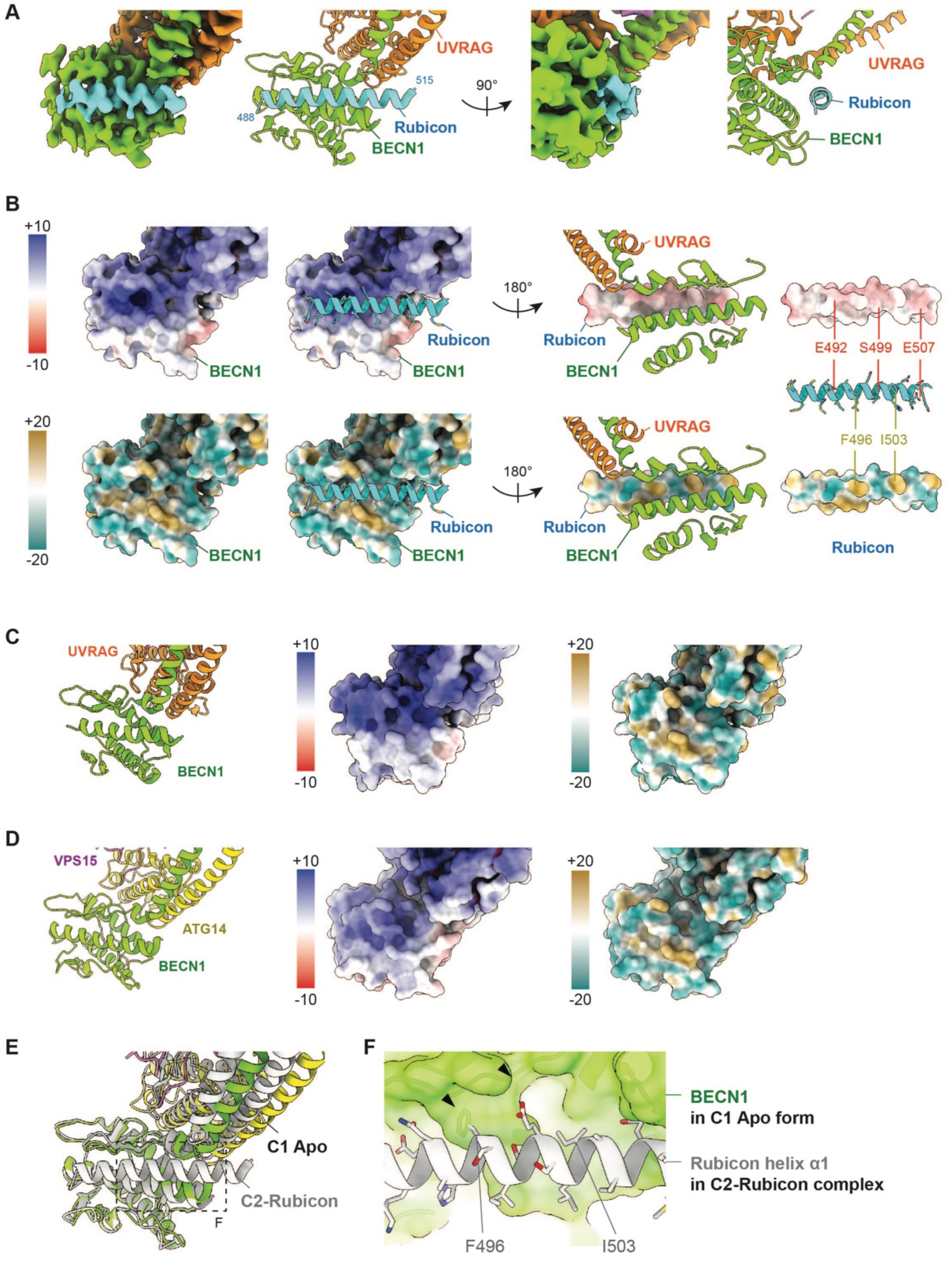
Structural analysis of the Rubicon–PI3KC3-C2 interaction. (A) Close-up views of the cryo-EM density (left) and the corresponding atomic model (right) of the Rubicon-binding site. (B) Electrostatic potential (red, −10; blue, +10) and molecular lipophilicity potential (cyan, −20; yellow, +20) surfaces of BECN1 and Rubicon. Residues selected for mutagenesis are indicated. (C and D) Corresponding views of the BECN1^BARA^ domain in the unbound PI3KC3-C2 (C) and PI3KC3- C1 (D) structures, showing the atomic model (left), electrostatic surface (middle), and molecular lipophilicity potential surface (right). (E) Superposition of the Rubicon complex (gray) onto the unbound PI3KC3-C1 structure following alignment of the BECN1^BARA^ domains. (F) Close-up view of the Rubicon-binding site in the unbound PI3KC3-C1 structure. BECN1 is shown as a transparent surface (green), and Rubicon is shown as ribbons and sticks (gray). The steric clash is indicated by black arrows.

**Fig. S3.**
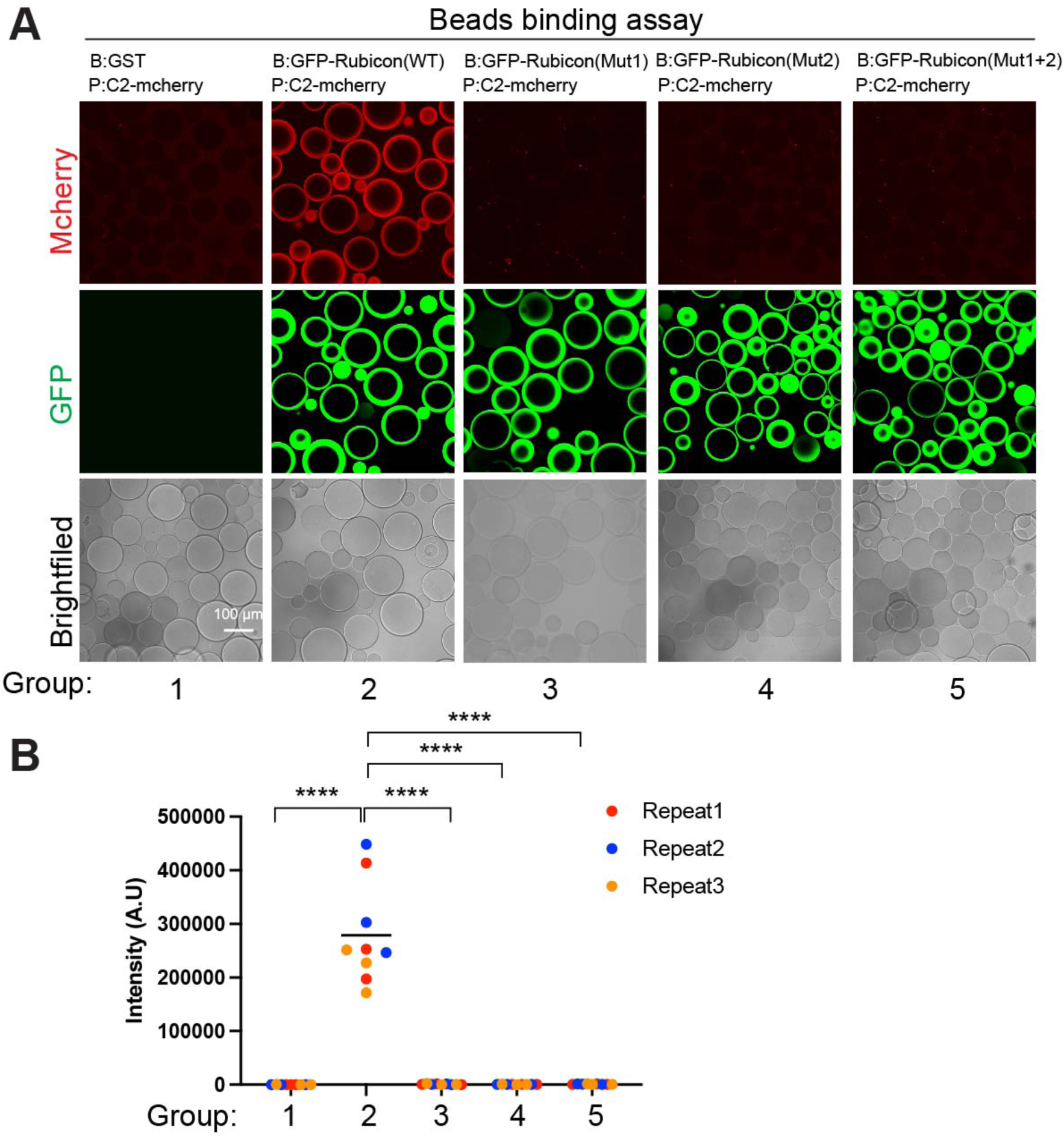
Complementary bead-binding assay. (A) Complementary GST bead-binding assay corresponding to that shown in Fig. 2A, with the bait and prey proteins exchanged. GSH beads coated with GST- and GFP-tagged WT Rubicon fragments (478-512) or the indicated mutants were incubated with mCherry-tagged PI3KC3-C2 or PI3KC3-C1. (B) Quantification of the GST bead-binding assay. Statistical significance was determined by one-way ANOVA followed by Bonferroni’s multiple-comparison test. Each differently colored dot represents an independent biological replicate, and each biological replicate includes 3 technical replicates. The average fluorescence intensity of all beads in each image was calculated. A.U represents arbitrary units. ****P < 0.0001 and ns: not significant. Data are presented as mean ± s.d.

**Fig. S4.**
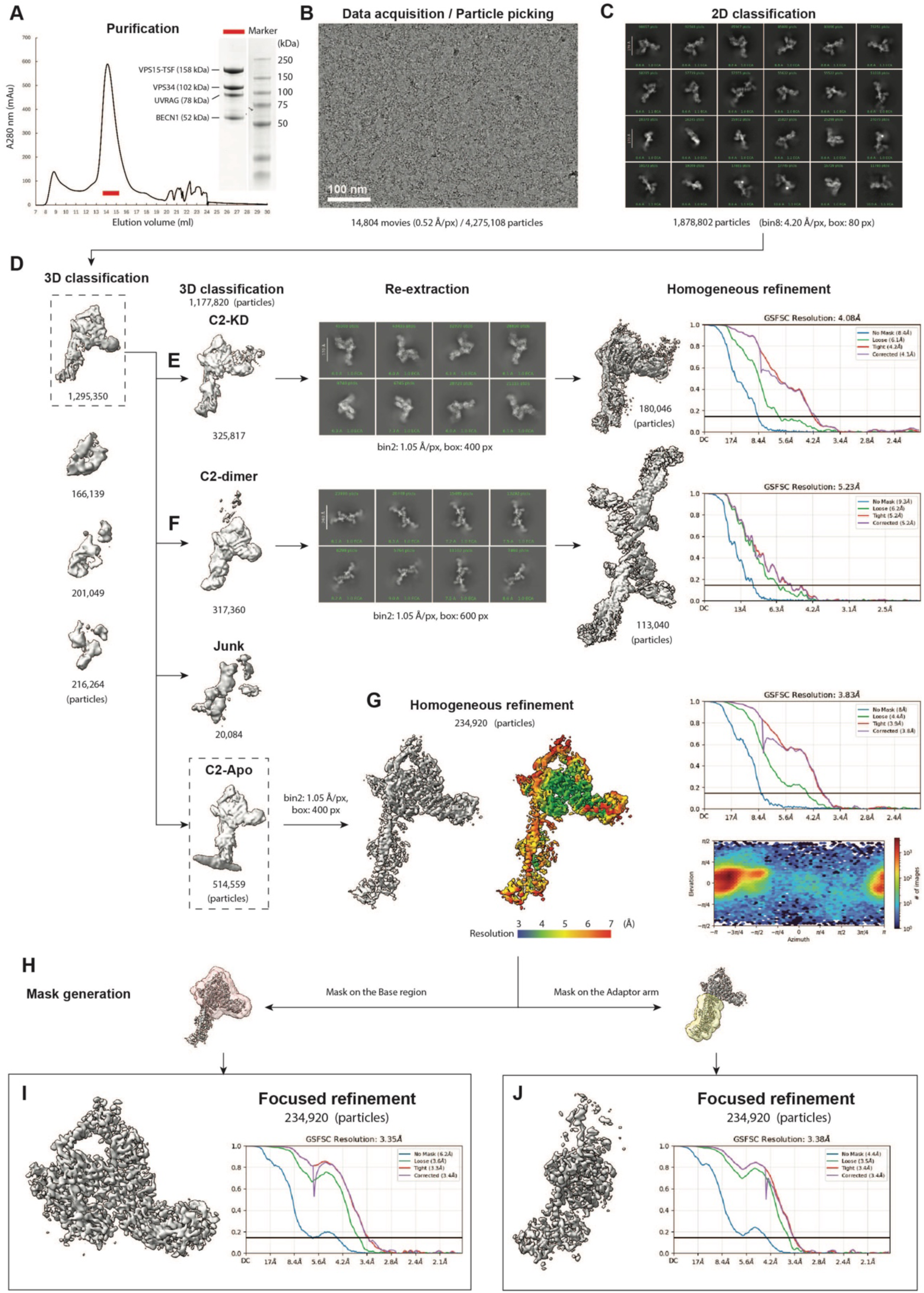
Cryo-EM sample preparation, image acquisition, and data processing of the unbound PI3KC3-C2. (A) Size-exclusion chromatography (SEC) profile of the unbound PI3KC3- C2 complex. The inset shows an SDS–PAGE analysis of the peak fraction (red bar). (B) Representative cryo-EM micrograph. (C) Representative 2D class averages. (D) Results of the initial round of 3D classification. (E) Classification and map reconstruction of PI3KC3-C2 particles containing well-resolved VPS34 kinase domain density. (F) Classification and map reconstruction of dimeric PI3KC3-C2 particles. (G) Consensus refinement of the final particle stack, with the corresponding FSC curve and local resolution map. (H) Mask generation for focused refinement. (I and J) Results of focused refinement, with the corresponding cryo-EM maps and FSC curves.

**Fig. S5.**
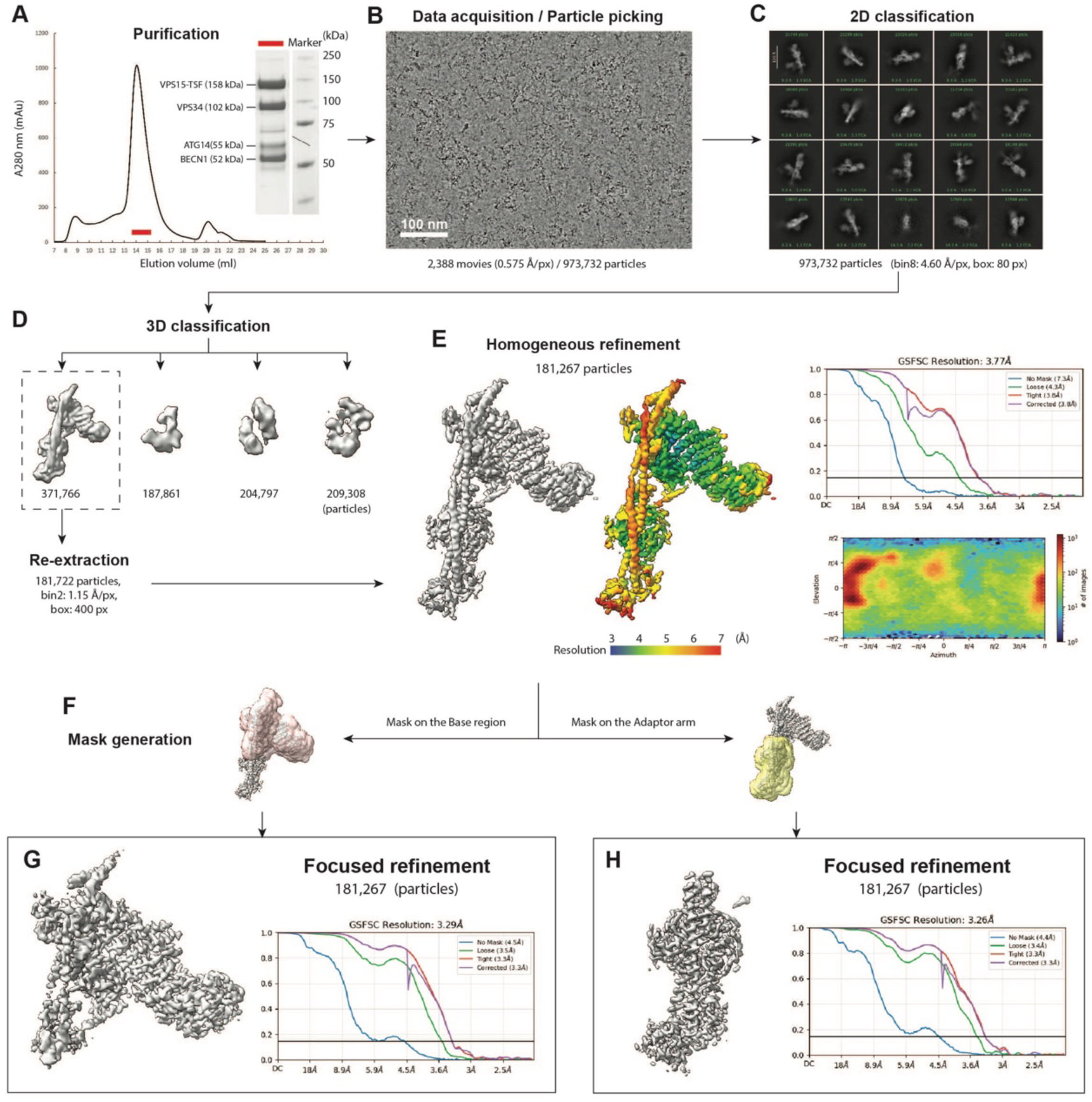
Cryo-EM sample preparation, image acquisition, and data processing of the unbound PI3KC3-C1. (A) Size-exclusion chromatography (SEC) profile of the unbound PI3KC3- C1 complex. The inset shows an SDS–PAGE analysis of the peak fraction (red bar). (B) Representative cryo-EM micrograph. (C) Representative 2D class averages. (D) Results of the initial round of 3D classification. (E) Consensus refinement of the final particle stack, with the corresponding FSC curve and local resolution map. (F) Mask generation for focused refinement. (G and H) Results of focused refinement, with the corresponding cryo-EM maps and FSC curves.

**Fig. S6.**
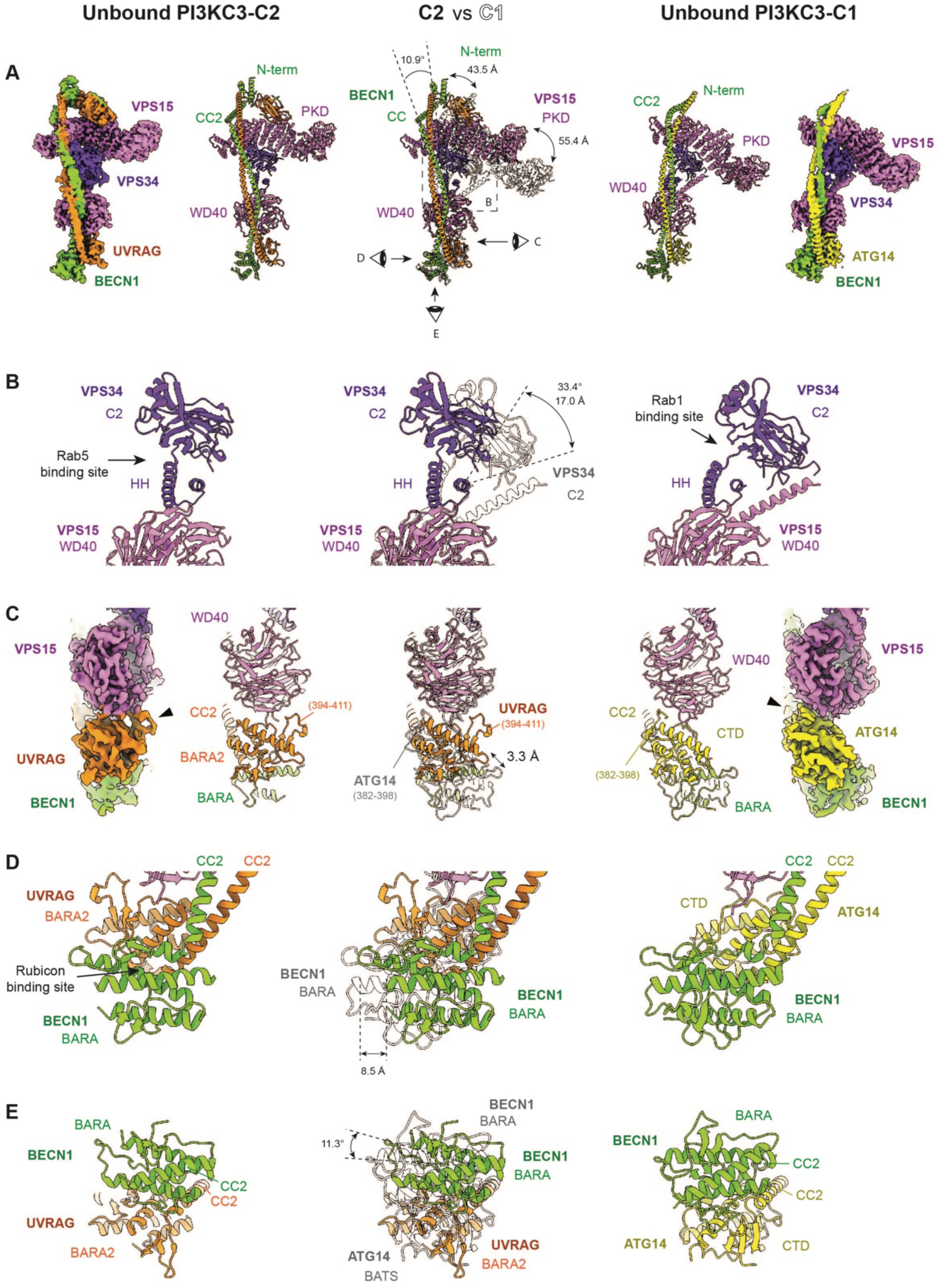
Conformational differences between the unbound PI3KC3-C2 and PI3KC3-C1 complexes. (A) Overall cryo-EM maps and corresponding atomic models of the unbound PI3KC3-C2 (left) and PI3KC3-C1 (right) complexes. Subunits and domains are labeled. The superposition of the two complexes following alignment of the VPS15^WD40^ domains is shown in the center, with the unbound PI3KC3-C1 structure displayed as a transparent overlay. Representative translations, rotations, and viewing directions for the subsequent panels are indicated. (B) Close-up views of the VPS34^C2^ domain. (C) Close-up views of the adaptor arm. (D and E) Close-up views of the BECN1^BARA^ domain.

**Table S1.** Cryo-EM data collection, refinement, and validation statistics.

|  | **Rubicon:**<br>**PI3KC3-C2** | **Unbound**<br>**PI3KC3-C2** | **Unbound**<br>**PI3KC3-C1** |
| --- | --- | --- | --- |
| EMDBID | EMD-74527 | EMD-74526 | EMD-76934 |
| PDBID | 9ZPD | 9ZPC | 13BV |
| **Data collection and processing** |  |  |  |
| Magnification | 81,000 | 81,000 | 36,000 |
| Voltage (kV) | 300 | 300 | 200 |
| Electron exposure (e<sup>-</sup>/Å<sup>2</sup>) | 50 | 50 | 50 |
| Defocus range (μm) | -0.8 to -2.0 | -0.8 to -2.0 | -0.8 to -2.0 |
| Physical pixel size (Å) | 0.940 | 1.050 | 1.115 |
| Symmetry imposed | C1 | C1 | C1 |
| Images | 21,885 | 14,804 | 2,338 |
| Initial particle images (no.) | 9,522,167 | 1,966,543 | 973,732 |
| Final particle images (no.) | 175,726 | 234,920 | 181,267 |
| Map resolution, Global (Å) | 3.38 | 3.83 | 3.77 |
| FSC threshold | 0.143 | 0.143 | 0.143 |
| **Refinement** |  |  |  |
| Map sharpening B factor (Å<sup>2</sup>) | 85.2 | 67.6 | 96.5 |
| Model composition |  |  |  |
| Non-hydrogen atoms | 17,405 | 16,338 | 15,991 |
| Protein residues | 2,149 | 2,016 | 1,976 |
| Ligands | 2 | 3 | 2 |
| B factors (Å<sup>2</sup>) |  |  |  |
| Protein | 95.01 | 78.51 | 74.59 |
| Ligands | 101.78 | 90.31 | 87.77 |
| R.m.s. deviations |  |  |  |
| Bond lengths (Å) | 0.004 | 0.008 | 0.004 |
| Bond angles (°) | 0.629 | 0.933 | 0.611 |
| Validation |  |  |  |
| MolProbity score | 1.67 | 2.32 | 1.87 |
| Clashscore | 7.03 | 25.84 | 11.95 |
| Poor rotamers (%) | 0.10 | 0.06 | 0 |
| Ramachandran plot |  |  |  |
| Favored (%) | 95.93 | 93.67 | 95.89 |
| Allowed (%) | 4.07 | 6.33 | 4.11 |
| Disallowed (%) | 0 | 0 | 0 |

Movie S1 (separate file). Structural basis for the PI3KC3-C2-specific interaction with Rubicon.

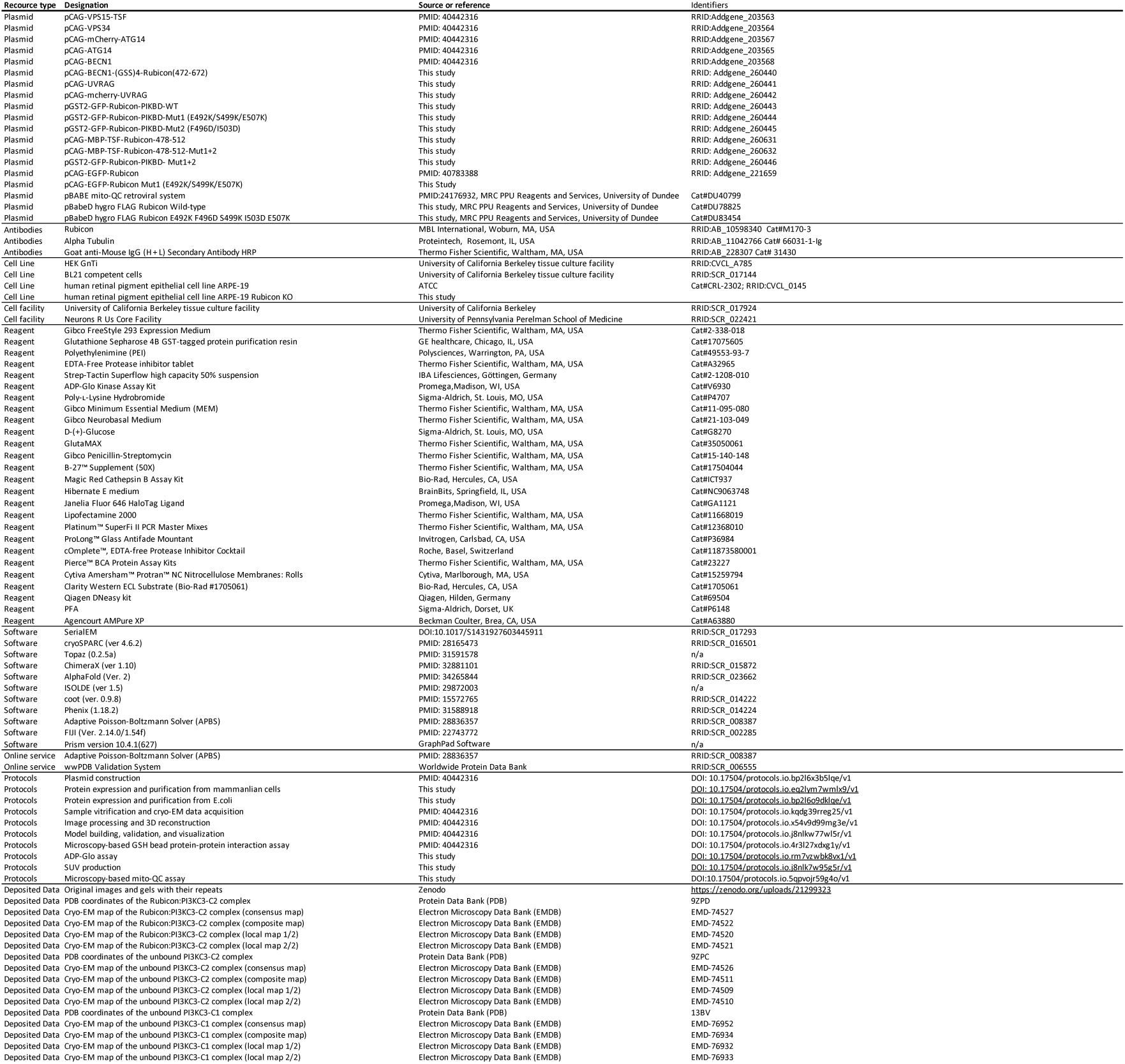

